# In Situ Inference of Copy Number Variations in Image-Based Spatial Transcriptomics

**DOI:** 10.1101/2025.07.02.662761

**Authors:** Augusta Jensen, Helena L. Crowell, Anna Pascual Reguant, Irene Ruano, Sabine Tejpar, Holger Heyn, Mats Nilsson, Sergio Marco Salas

## Abstract

Copy number variations (CNVs) drive cancer progression. So far, spatial CNV inference has relied on whole transcriptome-based sequencing technologies. However, advances in image-based spatial transcriptomics (iST) now enable high-plex gene measurement in situ. Here, we introduce an approach that adapts CNV inference to iST data, enabling spatial mapping of malignant clones and the tumor microenvironment, at single-cell resolution. Additionally, we assess how panel size and detection efficiency influence CNV inference.

## Introduction

Copy number variations (CNVs) are structural changes in the genome characterized by gains or losses of DNA segments, spanning from kilobases to whole chromosomes. Somatic CNVs are key contributors to cancer progression and are also valuable as biomarkers for diagnosis, prognosis, and therapy response^1–3^.

Since CNVs significantly impact gene expression^4^, multiple methods have been developed to infer the presence of CNVs in individual cells from transcriptomics data^5–7^, with variable success^8^. More recently, spatial transcriptomics (ST)^9^ technologies have enabled spatially resolved CNV inference, with all currently available methods operating on sequencing-based ST (sST). Examples of this include SpatialInferCNV^10^, which adapts inferCNV^6^ to work on sST data; STARCH^11^, which leverages each spot’s spatial position via a hidden Markov random field to boost CNV inference accuracy; and STmut^12^, which unifies point mutation, allelic imbalance, and CNV detection for comprehensive somatic alteration mapping. Although these tools have advanced our understanding of tumor evolution *in situ*^*13*^, most current and emerging sST technologies^14–17^ still lack the resolution and detection efficiency needed for accurate single-cell analysis of tumor clonality.

A promising alternative to spatially resolve tumor clonality at single-cell resolution could be to use imaging-based ST (iST) technologies (e.g. MERFISH^18^, CosMX^19^, Xenium^20^). However, their limited gene panels have been an obstacle for CNV inference. New iST technologies have recently boosted gene panel sizes, profiling thousands of genes in a single experiment (e.g., Xenium prime^21^ and CosMX Whole Transcriptome (WTx)^22^). Motivated by this enhanced gene coverage, we hypothesized that iST data could be used for single cell-resolved spatial CNV inference.

In this work, we present an approach for studying spatially resolved tumor clonality at single-cell resolution using CNV inference, and systematically evaluate the key technical factors that influence its performance.

## Results

To investigate the feasibility and accuracy of spatially resolved CNV inference in solid tumors, we designed a paired-section experimental setup integrating both spatial and dissociation-based transcriptomic profiling. We applied inferCNV to consecutive sections of a colorectal carcinoma (CRC) tumor^23^. One section was profiled by the iST CosMx SMI platform with the recently developed whole-transcriptome (WTx) panel^22^,complemented by histopathology annotations. The consecutive section was analyzed with single nuclei PATHO-seq (snPATHO-seq)^24^, a dissociation-based approach, and served as a single-cell reference for performance benchmarking/comparison/validation.

To enhance CNV detection, we implemented an RNA velocity-inspired smoothing step that averages each cell’s expression profile with those of its transcriptomic neighbors, reducing sparsity and improving signal-to-noise ratio (Methods). Smoothing substantially improved CNV detection in both datasets (Extended Data Fig. 1). Inferred CNVs were highly consistent between CosMx and snPATHO-seq, both identifying several CRC-associated alterations^25^, including gains in 13q and 20q and losses in 8p and 14q (Fig. 1a, Extended Data Fig. 2a).

**Figure 1:**
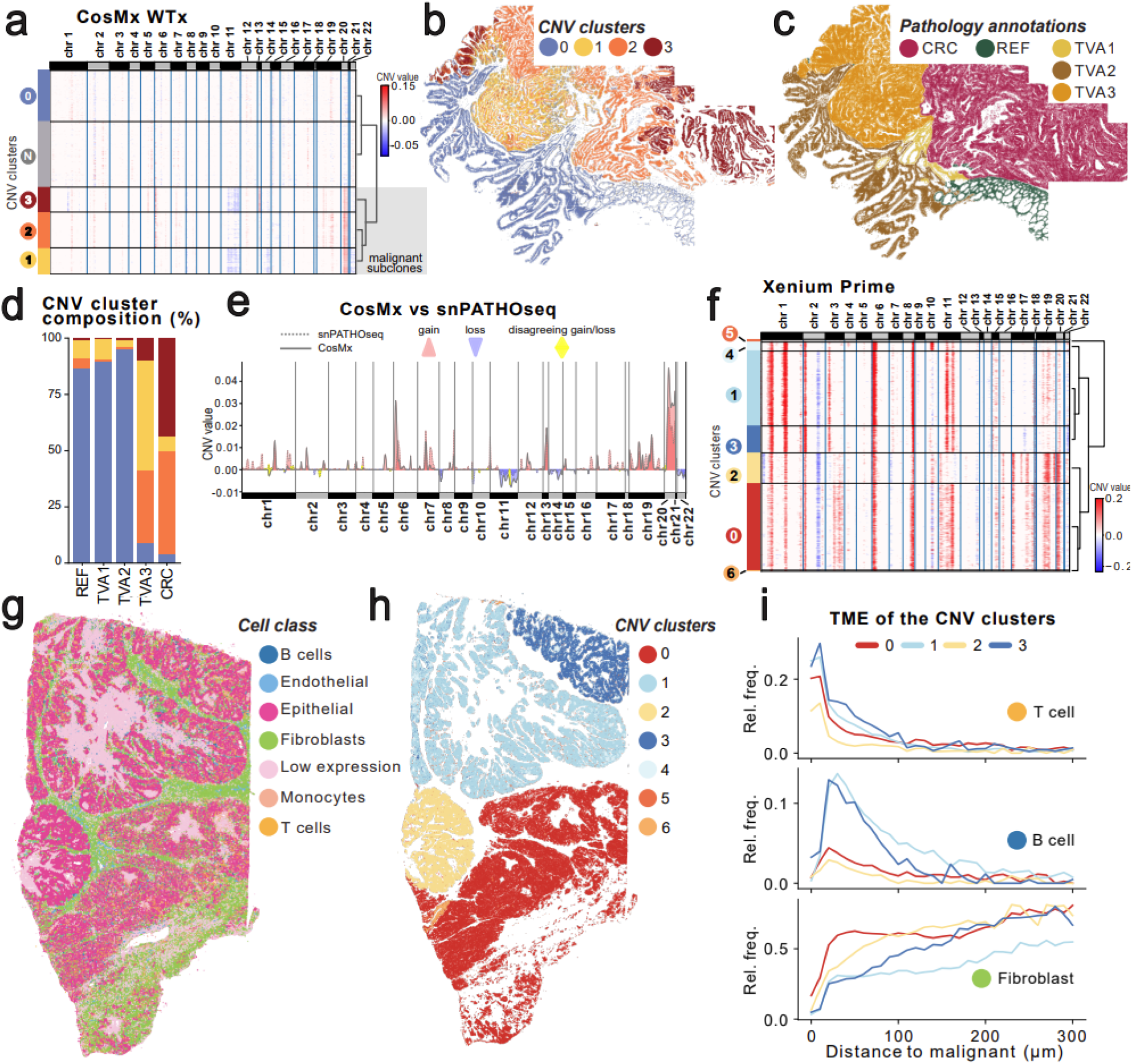
CNV inference and tumor subclone identification in two iST platforms. **a**, CNV heatmap of the colorectal carcinoma CosMx sample, showing gains (red) and losses (blue), including epithelial cells grouped by CNV cluster (0-3), and non-epithelial cells (N). Malignant subclones are marked in the dendrogram. **b**, Spatial map of epithelial CNV clusters presented in a. **c**, Histopathological annotations of the CosMx section, identifying healthy reference-like mucosa (REF), tubulovillous adenomas (TVA) stage 1-3, and cancerous colorectal carcinoma (CRC) regions. **d**, CNV cluster composition across histopathological compartments. **e**, Genome-wide comparison of average CNV profiles for the malignant CosMx (solid line) and snPATHO-seq (dotted line) cells, highlighting common gains (red), losses (blue) and disagreement between the datasets (yellow). **f**, CNV heatmap of the HGOC sample, illustrating inferred gains (red) and losses (blue). **g**, Map of cell classes in the HGOC dataset. **h**, Map of epithelial CNV clusters defined in f.. **i**, Relative frequencies of T cells, B cells, and fibroblasts plotted as a function of their distance to the nearest malignant subclone, as defined in h.

Focusing on the CosMx dataset, we identified four CNV-defined epithelial cell clusters (Fig. 1a). While one cluster lacked clear CNVs and grouped together with the non-epithelial cells, the other three exhibited prevalent CNVs, suggesting three malignant subclones. Their spatial distribution corresponded well with histopathological annotations^23^, with two dominating the annotated CRC region and a third enriched in late-stage premalignant tubulovillous adenoma (TVA3), suggesting early tumor progression. In contrast, the healthy mucosa (REF) and early-stage TVA regions were mainly composed of the CNV cluster lacking aberrant CNVs (Fig. 1b-d). Importantly, the three malignant CosMx subclones closely matched three CNV clusters identified by snPATHO-seq (Extended Data Fig. 2b, Methods), with their average CNV profiles displaying consistent patterns of gains and losses (Fig. 1e), thereby supporting the robustness of the approach.

Since most spatial platforms still rely on targeted panels with limited gene coverage^26^, we next assessed the broader applicability of our method by analyzing a second tumor type, high-grade serous ovarian carcinoma (HGSOC), profiled using a different iST platform, Xenium Prime (∼5000 genes). After annotating seven cell types (Extended Data Fig. 1a-b), we applied expression smoothing followed by CNV inference. Consistent chromosomal gains could be identified across epithelial cells, including 8q24 amplification, commonly found in ovarian cancer^27^ (Fig. 1f). The analysis of a control Xenium dataset from healthy lymph nodes, resulted in no detectable CNVs (Extended Data Fig. 3e-g), suggesting that HGSOC alterations reflect real biological events.

We next explored the spatial relationship between inferred CNV subclones and their surrounding tumor microenvironment in the HGSOC dataset. CNV profile clustering of epithelial cells revealed seven CNV clusters (Fig. 1f), with four of them (clusters 0-3) representing 95.7% of the total cells and exhibiting spatial compartmentalization (Fig. 1h). Notably, related subclones, as indicated by the CNV heatmap dendrogram (Fig. 1f), were spatially co-localized and associated with a more similar TME (Fig. 1i, Extended Data Fig. 3b, Methods). T-cells and B-cells were particularly enriched near subclones 1 and 3, whereas subclones 0 and 2 were situated in a fibroblast-rich region, linked to early relapse and progression^28^ (Fig. 1f).

While inferring CNVs in iST data is feasible, it’s limited by low transcript counts and gene panel size. To evaluate this, we simulated CNV subclones, applied key technical constraints of iST, and assessed CNV inference performance (Fig. 2a, Methods). As expected, reduced detection efficiency and gene panel size led to decreased CNV prediction and subclone identification accuracy (Fig. 2b). We computed metric scores for CNV gain and loss prediction, with CNV gains better inferred above 2,000 counts/cell, whereas CNV losses were poorly predicted even with high efficiency (AUC<0.8) (Fig. 2c). Interestingly, CNV gain accuracy improved with higher detection efficiency, whereas CNV losses showed a different trend, where datasets with fewer counts per cell were favored by a smaller gene panel. The performance of CNV gain prediction was saturated at around 1000 counts/cell, indicating diminishing returns beyond this point.

**Figure 2:**
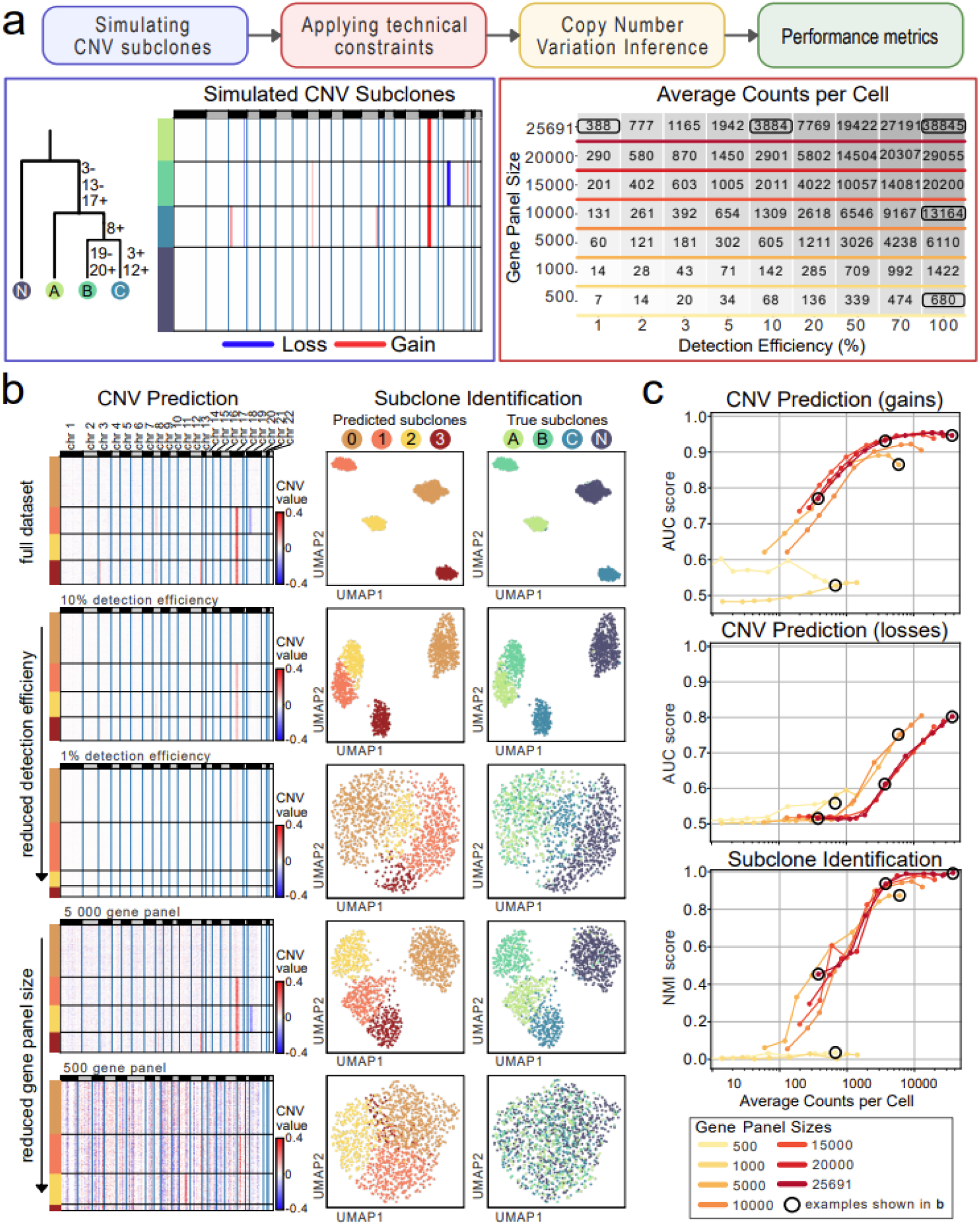
Evaluation of technical constraints affecting CNV inference. **a**, Workflow for evaluating how technical constraints affect CNV inference performance. Three CNV subclones (A, B, and C) were simulated alongside a normal (N) cell population lacking CNVs. The phylogenetic tree shows the gains and losses defining each subclone, which are also represented in the CNV heatmap, where gain (red) and losses (blue) reflect the simulated ground truth. Technical limitations of gene panel sizes and detection efficiency were applied to generate 63 unique datasets, five of which are shown in b. **b**, CNV heatmaps (left), shows inference results of the five selected datasets, two of which illustrate reduced detection efficiency, and two illustrate reduced gene panels. UMAPs (right) showing distributions of cells based on their inferred CNV profiles and compares their predicted vs true subclone identities. **c**, Performance metrics for CNV gain and loss prediction, as well as subclone identification. Black-circled data points highlight the five datasets shown in panel b.

Subclone identification was more influenced by efficiency than panel size. However, larger panels, especially the largest, delivered better overall results. Gene panels of 500 and 1,000 genes performed near random CNV prediction, and as expected, also failed in subclone identification. It is important to consider that CNV structure also influences inference performance. When repeating the analysis using different CNV sizes, larger CNVs were found to improve both CNV prediction and subclone identification (Extended Data Fig. 2).

## Discussion

Our study introduces a strategy to enable the inference of CNVs from imaging-based spatial transcriptomics (iST) data. This method broadens the utility of iST by enabling a new approach of subclone detection, while offering a major advantage over sequencing-based approaches: the ability to jointly analyze malignant subclones and their spatial microenvironment at single-cell resolution. In the HGSOC dataset, this enabled us to observe subclonal co-localization with distinct tumor microenvironment (TME) features, supporting the concept of tumor–TME co-evolution.

Our technical evaluation further clarified the limitations and requirements of CNV inference from iST data. Datasets with small gene panels (500 and 1000 genes) failed to produce meaningful CNV calls or subclone resolution, indicating that low-coverage panels are insufficient for this application. Detection efficiency also had an important impact on performance, where increases in transcript counts per cell improved CNV gain prediction and subclone identification. However, subclone identification performance plateaued around 1000 counts per cell and beyond this threshold, further increases offered diminishing returns.

We also found that subclones carrying larger CNVs were more reliably detected. This highlights that iST-based CNV inference is particularly effective for identifying dominant subclones with broad genomic alterations, but may lack the resolution to capture finer-scale heterogeneity. The quality of the input dataset plays a critical role in performance, and both platforms explored in this study (CosMx WTx and Xenium prime) exhibit data characteristics that lie near sharp performance thresholds. Notably, even small improvements in detection efficiency of imaging-based technologies could lead to substantial improvements in CNV inference accuracy and the robustness of subclone identification.

In conclusion, our study demonstrates that copy number variations can be directly inferred from in situ transcriptomics (iST) data by applying a transcriptomic-neighborhood smoothing approach to enhance signal detection. By applying and adapting in situ CNV inference across technologies and tissues, we reveal biologically relevant patterns and identify technical constraints. These findings open up new possibilities for studying somatic variation and tissue heterogeneity in situ. Future improvements in resolution and integration with multi-omics data will further enhance the power of this approach.

## Methods

### Gene expression smoothing

While initial CNV inference without smoothing revealed spatial alignment between two CosMx CNV clusters (showing gain in chr 20q) and the histopathological regions identified as cancerous colorectal carcinoma (CRC) and late-stage tubulovillous adenoma (TVA3), no obvious CNV profiles were generated in either of the two datasets (Extended Data Fig. 1; Fig. 1c).

We hypothesized that the weak CNV signals was likely caused by the inherently sparse transcript detection in both platforms^22,24^ (CosMx: 973 counts/cell, snPATHO-seq: 3,226 counts/cell, whereas a good scRNAseq dataset could generate more than 20,000 counts/cell^29^). Therefore, we applied a smoothing procedure that leverages the relationships between neighboring cells in a gene expression-based high-dimensional space, before inferring copy number variations (CNVs) in spatial datasets. Using a precomputed nearest-neighbor graph, connectivities were calculated based on the specified metric (either connectivities or distances). Smoothing increases with more neighbors. For each cell *i*, the smoothed expression value *M*_*i*_ was computed as a weighted average of its neighbors’ normalized (scanpy.pp.normalize_total) expression values:

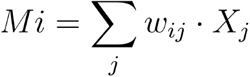

where *X*_*j*_ represents the normalized expression value for neighbor *j*, and *w*_*ij*_ is the weight derived from the connectivity matrix.

The smoothed data were stored in a new layer (*M*) within the AnnData object for downstream CNV analysis, including an additional normalization step (scanpy.pp.normalize_total) and log transformation (scanpy.pp.log1p). This step reduces noise in the raw data, enhances signal strength, and preserves spatial relationships, improving the reliability of CNV inference in single-cell and spatial datasets.

### CNV inference (by inferCNV)

To infer copy number variations (CNVs), we employed the python version of the inferCNV package (https://github.com/icbi-lab/infercnvpy). This was done on log-normalized or smoothed data, as described above. In summary, InferCNV calculates log fold changes (LFCs) across the genome for each cell, by subtracting the log-normalized gene expression of a predefined normal reference profile. To avoid false-positive CNVs caused by cell-type-specific gene expression (e.g., immunoglobulin or HLA genes), LFCs were adjusted in the following manner: (1) Genes with expression within the reference range were assigned an LFC of 0, (2) for genes expressed below the reference minimum, the minimum was subtracted and (3) for genes expressed above the reference maximum, the maximum was subtracted. The resulting LFCs were then smoothed along the genome using a defined window size and centered by subtracting the median expression of each cell. Noise was reduced by setting values below a dynamic threshold to zero, and a final median filter was applied to refine the signal, obtaining a matrix representing the inferred CNVs across cells and genomic regions. Specific parameters (window sizes, resolution and reference categories) used for each of the datasets processed can be found at https://github.com/Moldia/InSituCNV

Inferred CNV profiles from each dataset were visualized using functions from infercnvpy (infercnvpy.pl.chromosome_heatmap) as well as standard visualization tools from scanpy functions (scanpy.pl.umap, scanpy.pl.spatial), among others. To explore CNV-based cell subclonal structures, clustering was performed on the inferred CNV matrix obtained in previous steps. Following an analogous procedure of expression-based clustering in scanpy, we first computed k-nearest neighbors for each cell using the CNV profiles (infercnvpy.tl.neighbors). These neighbor graphs were then used to identify clusters via the Leiden algorithm (infercnvpy.tl.leiden) and low-dimensional representation of cells using UMAP (infercnvpy.tl.umap).

### CosMx and snPATHO-seq comparison

For consistency, both datasets were filtered to retain 14,055 shared genes. Gene expression was then log-normalized or smoothed, and CNV inference was carried out, as described in the sections above. After identifying subclones through clustering, we calculated cosine similarities between their average CNV profiles, revealing closely clustered subclones representing malignant cells. (Extended Data Fig. 2b). The average CNV profiles of these malignant cells were then plotted on top of each other, showing disagreeing regions (Fig. 1e)

### Tumor microenvironment analysis

Besides the identification and mapping of CNV clusters based on cells’ inferred CNV profile, in the HGOC Xenium sample analyzed, tumor microenvironment analysis was performed. Essentially, each cell previously identified as non-malignant by CNV analysis, was assigned to each closest malignant cell based on euclidean distances of their spatial positions. Then non-malignant cells were grouped based on the CNV cluster of their closest malignant cell and their cell type frequency was represented based on their distance to the closest malignant cell, obtaining cell type distribution profiles of each CNV clusters’ tumor microenvironment (Fig. 1i, Extended Data Fig. 3b).

### Generating CNV simulated datasets

To test how technical limitations such as *read depth* and *gene panel size* affect the inference of CNVs, a simulated dataset was generated using a 10X Chromium 10x 3’ v3 scRNAseq lung organoid dataset^30^. The dataset was filtered to only contain cells from one of the two donors. To get rid of noise or naturally occurring CNVs in the dataset, in preparation for simulating CNVs, one cluster was selected after running CNV inference.

Eight genes were selected to define the location and the type (gain/loss) of simulated CNV regions. The size of each CNV region was randomly sampled within a predefined interval. Using these parameters, a template was generated to annotate, for each gene, whether it fell within a CNV region and whether it was subject to a copy number gain or loss.

Based on this template, three simulated subclones were defined to represent different stages of malignant clonal evolution (Fig. 2a). Cells of a certain cell type were randomly assigned to one of the malignant subclones (1, 2, or 3), or as normal (N), together with the cells of the other cell types.

To simulate the effects of CNVs on gene expression, the original count matrix was modified according to the CNV status of each gene *g* and cell *i*. For every gene marked as a gain or loss in a given subclone, the raw count *x*_*ig*_ was adjusted using a randomly sampled scaling factor *ρ* [1, *α*], where *α* is the maximum amplitude of CNV effect. Depending on whether the gene was marked as a gain or a loss in the subclone template, the raw counts were adjusted accordingly:

- For CNV gains:

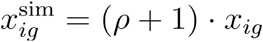
- For CNV losses:

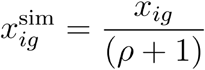

This approach maintained the natural variability in the data while introducing realistic, clone-specific CNV-driven expression changes.

Additionally two simulated datasets, with smaller (1–5 Mb) and larger (10-20 Mb) CNV regions, were generated to examine the impact of CNV structure beyond technical limitations (Extended Data Fig. 4).

### Modifying read depth and gene panel size

The simulated CNV datasets were made to mimic the technical constraints of detection efficiency and gene panel sizes. Datasets with varying read depths were generated by randomly downsampling (scanpy.pp.downsample_counts) the total number of counts in the count matrix to a fraction corresponding to 100%, 70%, 50%, 20%, 10%, 5%, 3%, 2% and 1% of the original CNV simulated dataset.

For each read-depth condition, we further reduced the *gene panel* by randomly removing genes, resulting in gene panel sizes of 20k, 15k, 10k, 5k, 1k, and 500 genes, as well as the full dataset gene panel, containing 25691 genes.

This process resulted in 63 distinct dataset variations (Fig. 2a), covering a range of detection efficiencies and gene panel sizes, each dataset with a unique amount of average counts per cell. The largest dataset retained the full 25691-gene panel with an average read depth of 37,548 counts per cell, whereas the most reduced dataset with a gene panel size of 500 genes had an average read depth of seven counts per cell.

### Quantitative metrics for CNV inference performance evaluation

Quantitative metrics were designed to evaluate the performance of CNV inference at different gene detection efficiencies and gene panel sizes. Essentially, two main tasks were quantitatively evaluated: CNV inference and subclone identification. For each of the tasks, specific metrics were employed.

#### CNV prediction metrics

To further evaluate how robustly and sensitively the CNV inference algorithm could detect the “true” CNVs in the data, specific metrics were employed to compare the inferred CNV patterns against the ground truth. **Receiver Operating Characteristic (ROC) Area Under the Curve (AUC)** and **F1 Score** were chosen as the primary performance metrics for this evaluation due to their relevance in assessing classification accuracy in the context of CNV detection.

First, ROC AUC measures the ability of the algorithm to distinguish between true positives and false positives across varying thresholds, providing a comprehensive evaluation of the classifier’s overall performance. The formula for ROC AUC is derived from the integration of the **True Positive Rate (***TPR*) (also known as sensitivity) and **False Positive Rate (***FPR*) over all possible thresholds:

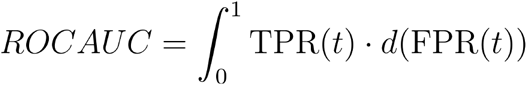

Where:

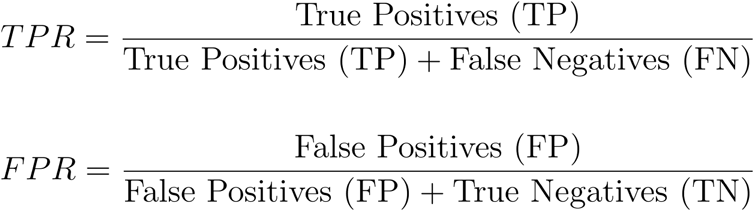

The ROC AUC ranges from 0 to 1, where a score of 1 indicates perfect classification, and 0.5 indicates performance no better than random guessing.

The **F1 Score** was calculated to provide a balanced measure of the algorithm’s precision and recall. Precision (*P*) measures the proportion of predicted CNVs that are correct, while recall (*R*) measures the proportion of true CNVs that were identified. The F1 Score is the harmonic mean of precision and recall, defined mathematically as:

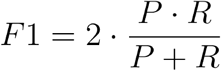

Where:

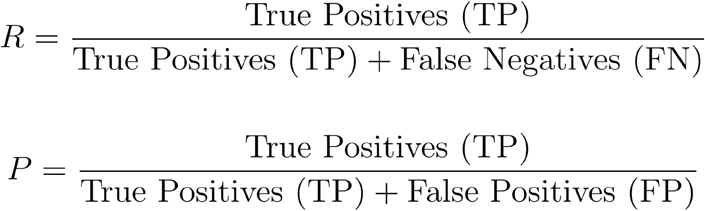

### Subclone identification metrics

To evaluate the performance of subclone identification, a similar approach was adopted using clustering comparison metrics. The Adjusted Rand Index (ARI) and Normalized Mutual Information (NMI) were computed to quantitatively assess the concordance between the predicted subclone clusters and the ground truth labels, derived from simulated data. These metrics were chosen because they provide complementary perspectives on clustering accuracy.

On the one hand, ARI evaluates the level of agreement between the two clustering strategies by considering all possible pairs of points and determining whether they are consistently assigned to the same or different clusters. It is adjusted for chance, ensuring that random cluster assignments yield an expected value close to zero. A perfect ARI score of 1 indicates that the computed clusters match the reference labels exactly.

On the other hand, NMI measures the shared information between the two clustering outcomes normalized by the average of their entropy. It is particularly useful for evaluating cluster consistency when the number of clusters differs between the reference and computed labels. An NMI score of 1 indicates that the two clusterings share complete mutual information, demonstrating perfect alignment.

## Data availability

The CosMx WTx and snPATHO-seq data from the CRC sample 221 used in this study are available from Crowell et al., “*Tracing colorectal malignancy transformation from cell to tissue scale*,” bioRxiv (2025) https://doi.org/10.1101/2025.06.23.660674. The 10X Xenium datasets used through the study are publicly available at the following links: https://www.10xgenomics.com/datasets/xenium-prime-fresh-frozen-human-ovary (high grade ovarian cancer - Xenium prime dataset) and https://www.10xgenomics.com/datasets/preview-data-xenium-prime-gene-expression (lymph node Xenium prime dataset). The 10X scRNA-seq dataset used for CNV simulation is available at https://cellxgene.cziscience.com/collections/e9cf4e8d-05ed-4d95-b550-af35ca219390.

## Code availability

All code used to perform the analysis discussed in this manuscript, and generate the figures presented, is accessible at https://github.com/Moldia/InSituCNV.

## Acknowledgements

M.N. is supported by the Swedish Research Council (project 2024-02533), Cancerfonden (project 24-3457), U-CAN, the Erling Persson Foundation, and the Horizon Europe Mission on Cancer “Spacetime” (project 101136552). H.L.C. acknowledges support by the Swiss National Science Foundation (project grant number 222136). A.P.R. has received funding from the MCIN/AEI/10.13039/501100011033 and FSE+ (RYC2022-035848-I) and the MICIU/AEI/10.13039/501100011033/ FEDER/UE (PID2023-148687OB-I00). H.H. has received funding from the European Union’s H2020 research and innovation program (848028), from the European Research Council (ERC) (810287), from the Ministerio de Ciencia e Innovación (MCI) (PID2020-115439GB-I00 and PLEC2021-007654), from LaCaixa Foundation (HR22-0031 and HR22-0172), from the Generalitat de Catalunya through the Suport Grups de Recerca AGAUR (2021-SGR) and from ERA-NET Neuron/Ministerio de Ciencia e Innovación (MCI) (PCI2022-133012). H.H. was supported by an ASPIRE Award from The Mark Foundation for Cancer Research and the Scientific Foundation of the Spanish Association Against Cancer.

## Extended Data

**Extended Data Figure 1:**
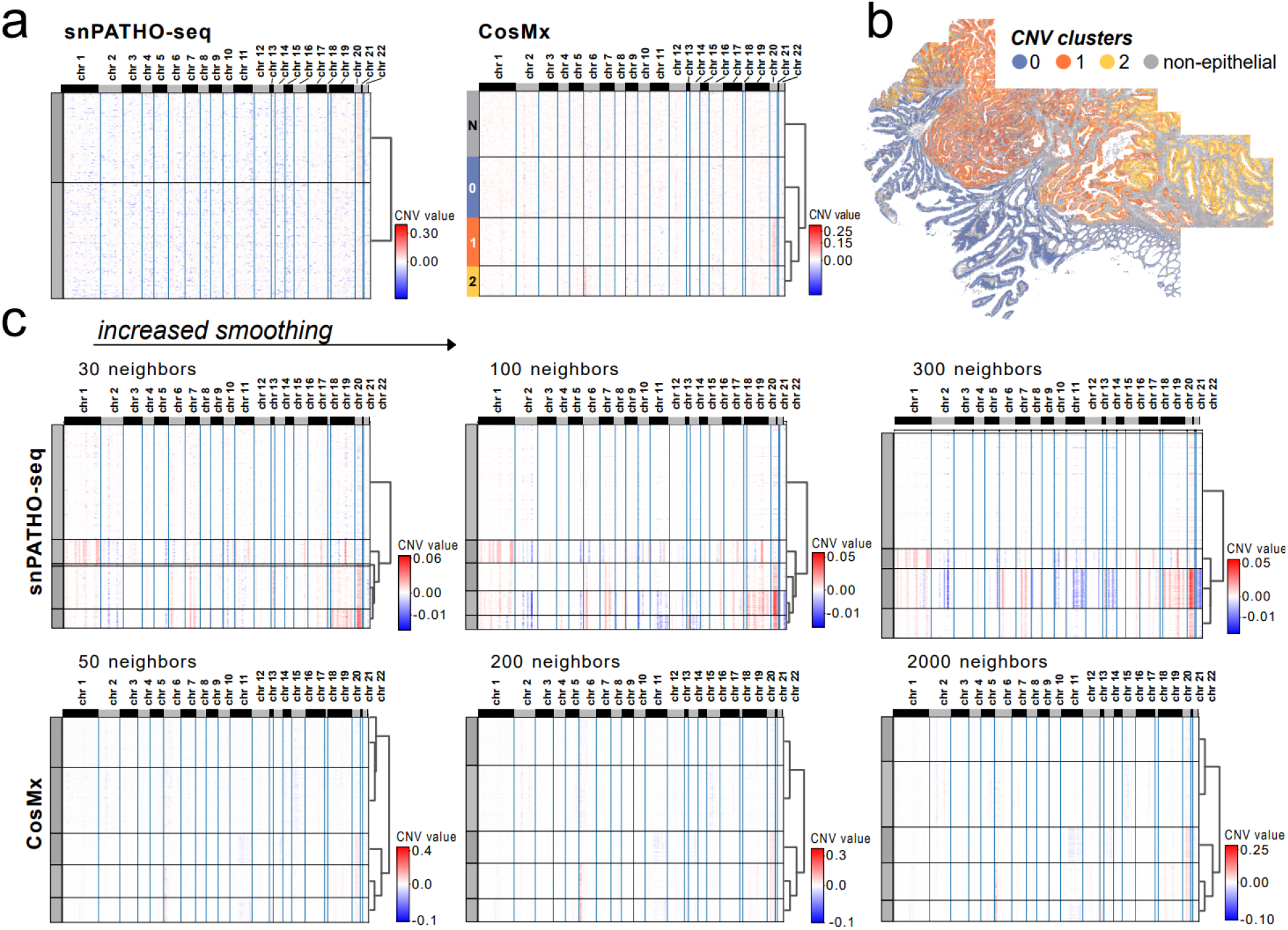
CNV inference with increased smoothing. **a**, CNV inference of snPATHO-seq and CosMx data without smoothing. CNV heatmaps display copy number gains in red and losses in blue. **b**, Map of CNV clusters identified in a. **c**, CNV heatmaps of snPATHO-seq and CosMx with increased smoothing, achieved by incorporating signals from additional neighboring cells to enhance CNV inference.

**Extended Data Figure 2:**
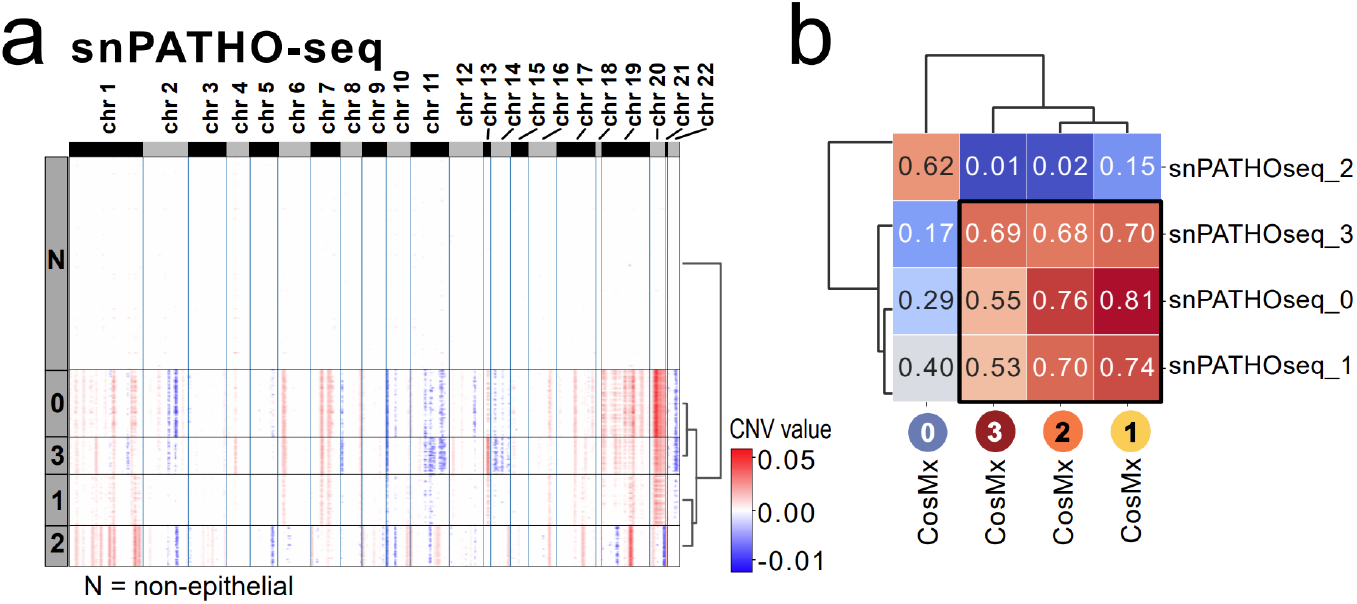
CNV profile comparison between CosMx and snPATHO-seq. **a**, CNV heatmap of the snPATHO-seq dataset, showing gains in red and losses in blue. **b**, Cosine similarity heatmap comparing CNV clusters from CosMx and snPATHO-seq. The black square highlights malignant subclones.

**Extended Data Figure 3:**
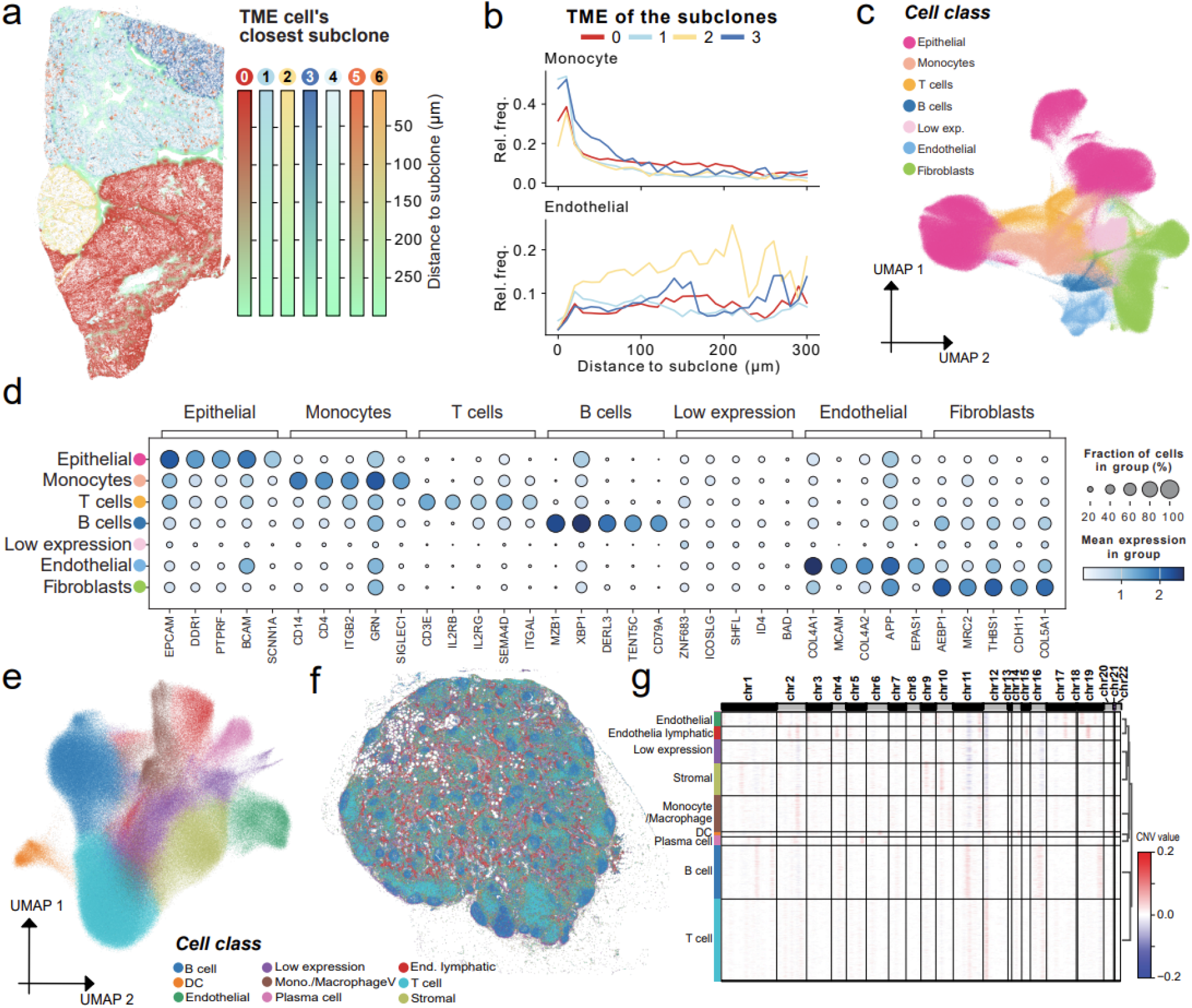
Supplementary analysis of Xenium Prime data. **a**, Map of HGSOC’s TME cells, with a color scale indicating each cell’s distance (um) to the nearest malignant subclone, as defined by CNV clusters in Fig. 1h. **b**, Relative frequencies of endothelial and monocyte cells, across the distance to each malignant subclone. **c**, UMAP of ovarian sample cells, colored by cell class. **d**, Expression of the top 5 differentially expressed genes by cell class. **e**, UMAP of the cells, colored by class in the lymph node sample. **f**, Spatial map of the cell classes in the lymph node. **g**, CNV heatmap of lymph node.

**Extended Data Fig. 4:**
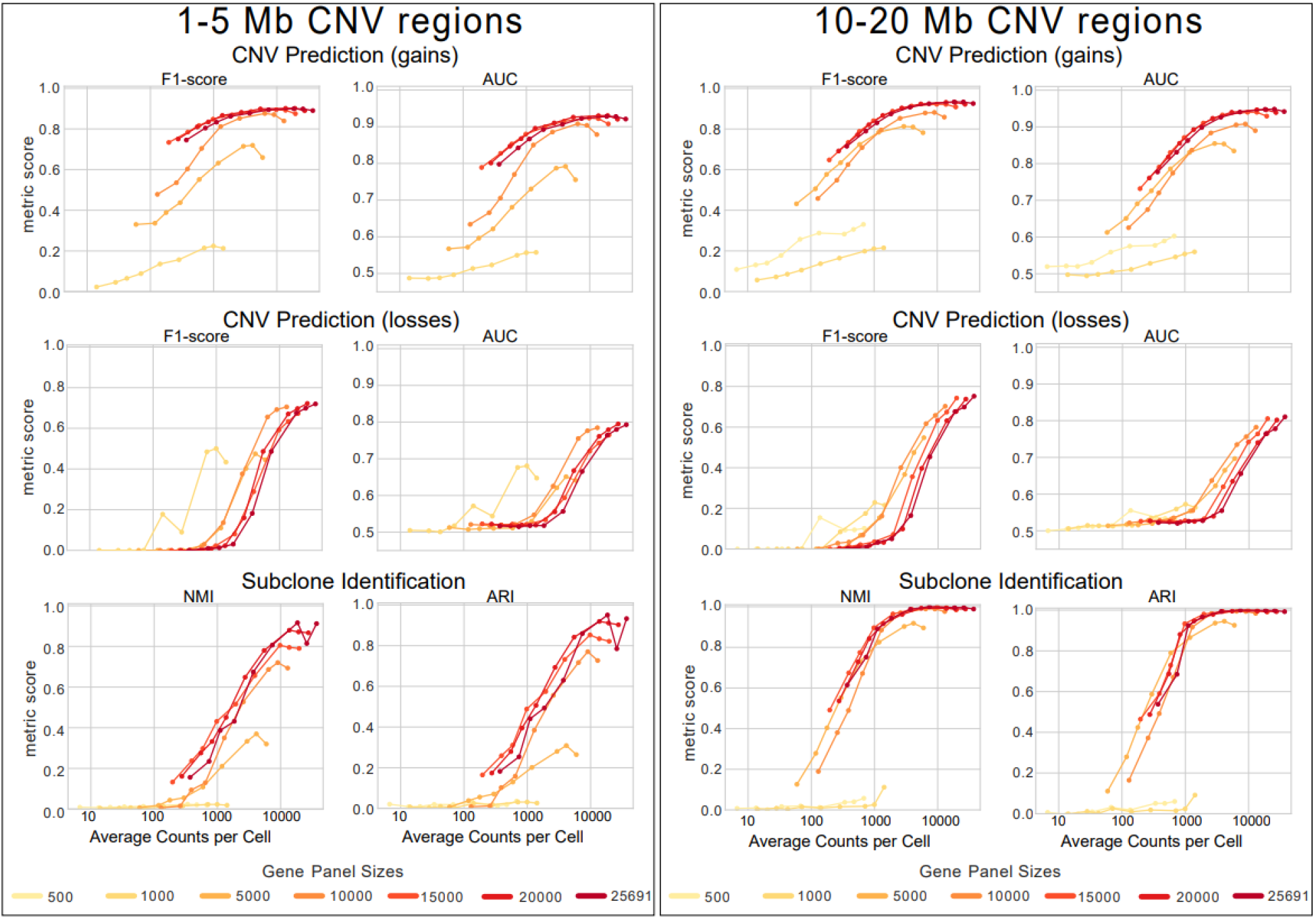
Performance metrics across different CNV region sizes. Performance metrics for CNV gain and loss prediction and subclone identification across 63 simulated datasets, featuring CNV regions of 1–5 Mb (left) and 10-20 Mb (right), and imposed technical constraints. Datasets were grouped by gene panel sizes, indicated by line color. X-axis represents average counts per cell with Y-axis representing performance metrics including: F1-score and Area Under the ROC Curve (AUC) for CNV prediction, and Adjusted Rand Index (ARI) and Normalized Mutual Information (NMI) for subclone identification. In some graphs, the 500-gene panel is absent due to missing ground truth values.

## References

1. Steele, C. D. et al. Signatures of copy number alterations in human cancer. Nature 606, 984–991 (2022).

2. Carey-Smith, S. L., Kotecha, R. S., Cheung, L. C. & Malinge, S. Insights into the Clinical, Biological and Therapeutic Impact of Copy Number Alteration in Cancer. Int. J. Mol. Sci. 25, 6815 (2024).

3. Redon, R. et al. Global variation in copy number in the human genome. Nature 444, 444–454 (2006).

4. Shao, X. et al. Copy number variation is highly correlated with differential gene expression: a pan-cancer study. BMC Méd. Genet. 20, 175 (2019).

5. Gao, R. et al. Delineating copy number and clonal substructure in human tumors from single-cell transcriptomes. Nat. Biotechnol. 39, 599–608 (2021).

6. Patel, A. P. et al. Single-cell RNA-seq highlights intratumoral heterogeneity in primary glioblastoma. Science 344, 1396–1401 (2014).

7. Falco, A. D., Caruso, F., Su, X.-D., Iavarone, A. & Ceccarelli, M. A variational algorithm to detect the clonal copy number substructure of tumors from scRNA-seq data. Nat. Commun. 14, 1074 (2023).

8. Chen, X. et al. A benchmarking study of copy number variation inference methods using single-cell RNA-sequencing data. bioRxiv 2024.09.09.612120 (2024) doi:10.1101/2024.09.09.612120.

9. Moses, L. & Pachter, L. Museum of spatial transcriptomics. Nat. Methods 19, 534–546 (2022).

10. Erickson, A. et al. Spatially resolved clonal copy number alterations in benign and malignant tissue. Nature 608, 360–367 (2022).

11. Elyanow, R., Zeira, R., Land, M. & Raphael, B. J. STARCH: copy number and clone inference from spatial transcriptomics data. Phys. Biol. 18, 035001 (2021).

12. Chen, L. et al. STmut: a framework for visualizing somatic alterations in spatial transcriptomics data of cancer. Genome Biol. 24, 273 (2023).

13. Mo, C.-K. et al. Tumour evolution and microenvironment interactions in 2D and 3D space. Nature 634, 1178–1186 (2024).

14. Marshall, J. L. et al. High-resolution Slide-seqV2 spatial transcriptomics enables discovery of disease-specific cell neighborhoods and pathways. iScience 25, 104097 (2022).

15. Chen, A. et al. Spatiotemporal transcriptomic atlas of mouse organogenesis using DNA nanoball-patterned arrays. Cell 185, 1777-1792.e21 (2022).

16. Oliveira, M. F. et al. Characterization of immune cell populations in the tumor microenvironment of colorectal cancer using high definition spatial profiling. bioRxiv 2024.06.04.597233 (2024) doi:10.1101/2024.06.04.597233.

17. Liu, Y. et al. High-Spatial-Resolution Multi-Omics Sequencing via Deterministic Barcoding in Tissue. Cell 183, 1665-1681.e18 (2020).

18. Chen, K. H., Boettiger, A. N., Moffitt, J. R., Wang, S. & Zhuang, X. Spatially resolved, highly multiplexed RNA profiling in single cells. Science 348, aaa6090 (2015).

19. He, S. et al. High-plex imaging of RNA and proteins at subcellular resolution in fixed tissue by spatial molecular imaging. Nat. Biotechnol. 40, 1794–1806 (2022).

20. Janesick, A. et al. High resolution mapping of the tumor microenvironment using integrated single-cell, spatial and in situ analysis. Nat. Commun. 14, 8353 (2023).

21. Ren, P. et al. Systematic Benchmarking of High-Throughput Subcellular Spatial Transcriptomics Platforms. bioRxiv 2024.12.23.630033 (2024) doi:10.1101/2024.12.23.630033.

22. Khafizov, R. et al. Sub-cellular Imaging of the Entire Protein-Coding Human Transcriptome (18933-plex) on FFPE Tissue Using Spatial Molecular Imaging. bioRxiv 2024.11.27.625536 (2024) doi:10.1101/2024.11.27.625536.

23. Crowell, H. L. et al. Tracing colorectal malignancy transformation from cell to tissue scale. bioRxiv 2025.06.23.660674 (2025) doi:10.1101/2025.06.23.660674.

24. Wang, T. et al. snPATHO-seq, a versatile FFPE single-nucleus RNA sequencing method to unlock pathology archives. Commun. Biol. 7, 1340 (2024).

25. Debattista, J., Grech, L., Scerri, C. & Grech, G. Copy Number Variations as Determinants of Colorectal Tumor Progression in Liquid Biopsies. Int. J. Mol. Sci. 24, 1738 (2023).

26. Jonghe, J. D. et al. scTrends: A living review of commercial single-cell and spatial ‘omic technologies. Cell Genom. 4, 100723 (2024).

27. Goode, E. L. et al. A genome-wide association study identifies susceptibility loci for ovarian cancer at 2q31 and 8q24. Nat. Genet. 42, 874–879 (2010).

28. Xu, A. M. et al. Spatiotemporal architecture of immune cells and cancer-associated fibroblasts in high-grade serous ovarian carcinoma. Sci. Adv. 10, eadk8805 (2024).

29. Yamawaki, T. M. et al. Systematic comparison of high-throughput single-cell RNA-seq methods for immune cell profiling. BMC Genom. 22, 66 (2021).

30. Kim, J. et al. Transcriptomic Analysis of Air–Liquid Interface Culture in Human Lung Organoids Reveals Regulators of Epithelial Differentiation. Cells 13, 1991 (2024).

